# Predator feeding rates may often be unsaturated under typical prey densities

**DOI:** 10.1101/2022.08.09.503207

**Authors:** Kyle E. Coblentz, Mark Novak, John P. DeLong

**Affiliations:** School of Biological Sciences, University of Nebraska-Lincoln, Lincoln, NE; Department of Integrative Biology, Oregon State University, Corvallis, OR

**Author notes:** Corresponding Author e, 402 Manter Hall, University of Nebraska-Lincoln, Lincoln, NE, USA 68588. Statement of Authorship: All authors designed the research, KEC performed the analyses with feedback from JPD and MN, KEC wrote the first draft of the manuscript, all authors contributed to revisions. Data Accessibility Statement: All data and code for the analysis of the data are available at https://github.com/KyleCoblentz/FeedingRateSaturation. This repository will be permanently archived on Zenodo upon acceptance.

**Keywords:** Predator-Prey Interactions, Functional Response, Nonlinearity, Interaction Strength, Consumer-Resource, Mass-Abundance Scaling

## Abstract

Predator feeding rates (described by their functional response) must saturate at high prey densities. Although thousands of manipulative functional response experiments show feeding rate saturation at high densities under controlled conditions, it is unclear *how* saturated feeding rates are at natural prey densities. The general degree of feeding rate saturation has important implications for the processes determining feeding rates and how they respond to changes in prey density. To address this, we linked two databases – one of functional response parameters and one on mass-abundance scaling – through prey mass to calculate a feeding rate saturation index. We find that: 1) feeding rates may commonly be unsaturated and 2) the degree of saturation varies with predator and prey taxonomic identities and body sizes, habitat, interaction dimension, and temperature. These results reshape our conceptualization of predator-prey interactions in nature and suggest new research on the ecological and evolutionary implications of unsaturated feeding rates.

## Introduction

Predator functional responses describe predator feeding rates as a function of prey density and are a central component of theory on consumer-resource interactions (Solomon 1949; Holling 1959; Murdoch *et al*. 2013). As pointed out by Holling (1959), functional responses should saturate with increasing prey density because the time it takes to process prey items, generally referred to as the handling time, limits feeding rates at high prey densities. Since its inception, the idea of functional response saturation has become a canonical component of predator-predator theory, shaping the way we conceptualize predator-prey interactions and dynamics (Rosenzweig & MacArthur 1963; McCann 2011; Murdoch *et al*. 2013).

Logic and thousands of experiments make it clear that feeding rates are saturating functions of prey densities (Holling 1959; Jeschke *et al*. 2004; Novak & Stouffer 2021; Uiterwaal *et al*. n.d.). However, it remains unclear *how* saturated feeding rates are under the prey densities that predators experience in nature. Some models and data suggest that, for carnivores, feeding rates should be saturated at the prey densities they experience (Jeschke 2007). This is because carnivores appear to be digestion-limited and satiated, or ‘full and lazy’ (Jeschke 2007). Indeed, some field functional response studies show saturated feeding rates over large ranges of prey densities (e.g. Messier 1994; Nielsen 1999; Gilg *et al*. 2006; Nilsen *et al*. 2009; Moustahfid *et al*. 2010; Moleón *et al*. 2012). However, other studies show saturated feeding rates at only one or a few observations at the highest observed prey densities (Angerbjörn 1989; Korpimaki & Norrdahl 1991; Redpath & Thirgood 1999; Sundell *et al*. 2000; Quinn *et al*. 2017; Coblentz *et al*. n.d.) or little to no evidence of feeding rate saturation (Novak 2010; Novak *et al*. 2017; Preston *et al*. 2018; Beardsell *et al*. 2021, n.d.; Coblentz *et al*. 2021). Therefore, it is currently unclear how saturated feeding rates are likely to be in general.

The saturation level of feeding rates has important implications for predator-prey interactions. On one extreme, if feeding rates are generally saturated at typical prey densities, then feeding rates are largely determined by handling times, will be near their maxima (the reciprocal of the handling time), and should show little response to changes in prey densities (Figure 1A). On the other extreme, if feeding rates are generally unsaturated, then feeding rates will largely be determined by space clearance rates (aka attack rates), will be lower than their potential maxima, and will dynamically respond to changes in prey densities (Figure 1B). Knowing whether predator feeding rates generally are saturated or unsaturated therefore would provide insights into the factors governing the strength of predator-prey interactions and how dynamic these interaction strengths may be.

**Figure 1.**
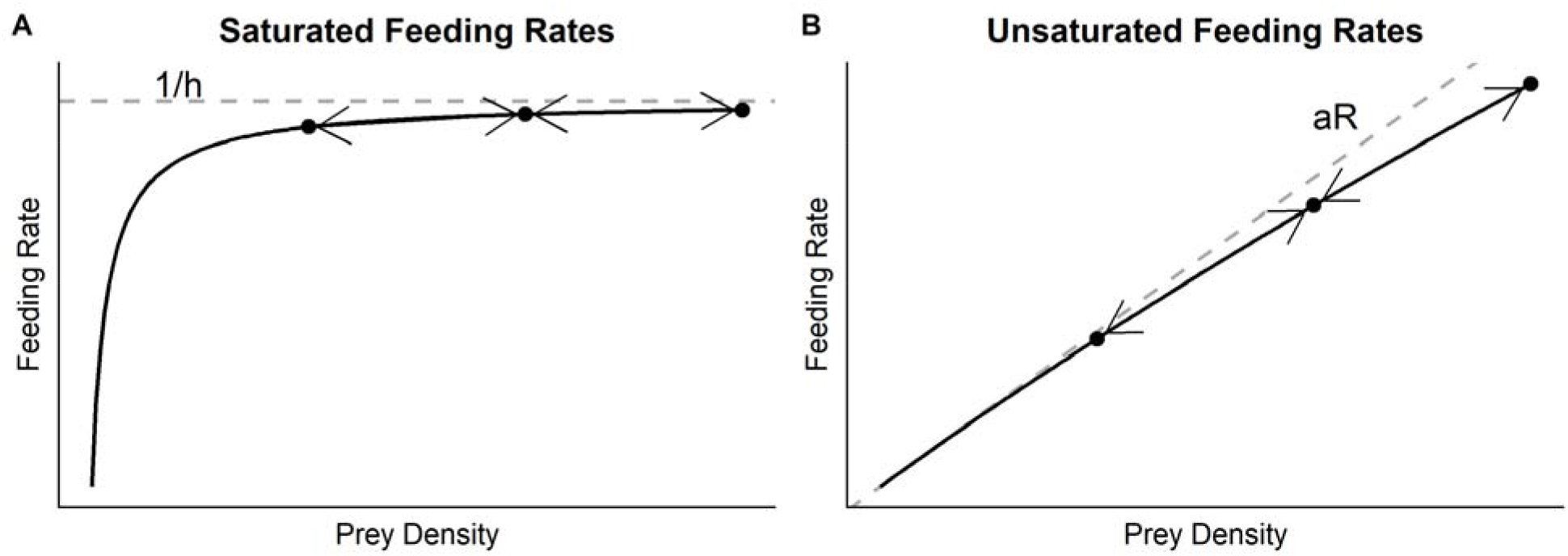
For saturated feeding rates (A), feeding rates are largely near their maximum as determined by the handling time (gray dashed line; 1/h) and will change little with changes in prey densities (represented by the dots and arrows). For unsaturated feeding rates (B), feeding rates are near the line determined by the space clearance or attack rate (a; gray dashed line) and will change close to proportionally with changes in prey densities (represented by the dots and arrows).

Here we combine two databases to investigate generalizations about how saturated feeding rates might be under typical prey field densities. The first database, the FoRAGE database (Functional Responses Across the Globe in all Ecosystems (Uiterwaal *et al*. n.d.)), contains estimates of saturating Type II functional response parameters from 2,598 functional response experiments, the vast majority of which are laboratory-based. Most these studies do not include field estimates of prey abundance. We therefore estimated prey abundances using a database on mass-abundance scaling (aka Damuth’s Rule (Damuth 1981; White *et al*. 2007)) containing 5,985 records of masses and field-estimated abundances across the major taxa on earth (Hatton et al. 2019). Combining the functional response parameters and prey masses from FoRAGE with estimates of prey field abundances from the mass-abundance scaling relationships, we estimate an index of feeding rate saturation to address two questions:

1. How saturated may predator feeding rates be under typical prey densities?
2. What covariates of a predator and prey’s biology or environmental context are related to the degree of feeding rate saturation?

Our results suggest that predator feeding rates are commonly unsaturated at prey densities experienced in the field. We also find that prey and predator taxonomic identity and body sizes, the dimensionality of their interaction, their habitat, and temperature explain a significant amount of the variation in feeding rate saturation.

## Materials and Methods

We first derive an index of feeding rate saturation. We then describe how we calculated the index using the FoRAGE database (Uiterwaal *et al*. n.d.) and data on mass-abundance scaling relationships (Hatton *et al*. 2019). Last, we describe our statistical analysis to examine how biological covariates influence the degree of feeding rate saturation.

### Feeding Rate Saturation Index

Our index of feeding rate saturation gives the proportional reduction in predator feeding rates due to saturation with increasing prey densities. For a predator with a saturating Type II functional response, we can derive the saturation index by comparing the feeding rates under the Type II functional response to the feeding rates under a non-saturating linear functional response. Under a linear (or Type I) functional response, the predator’s feeding rate is proportional to prey density R:

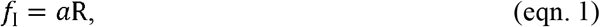

where *a* is the predator’s space clearance rate (Fig. 1B). Under a Holling Type II functional response, the predator’s feeding rate is

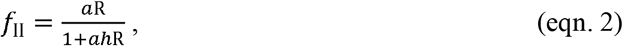

where, *h*, is the predator’s handling time. Our index of saturation *I* gives the proportional reduction in feeding rates between these two functional responses:

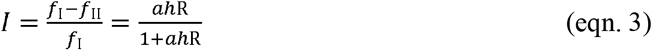

(see Supplemental Information S1 for a derivation). The saturation index can take values between zero and one. Values near zero indicate the feeding rate is relatively unsaturated and values near one indicate the feeding rate is close to complete saturation (i.e. maximum feeding rate). This index can also be generalized to many other common functional response forms (Supplemental Information S7).

### Estimating the Saturation Index

To estimate the saturation index, we need values for the space clearance rate (*a*), handling time (*h*), and prey density (R). We obtained estimates for the space clearance rates and handling times from the FoRAGE database (Uiterwaal *et al*. n.d.). The database also contains a suite of biological and contextual covariates that may influence the functional response parameters including the average mass of the predator and prey, the dimensionality of their interaction (2D, 3D, or 2.5D i.e. fractional between 2D and 3D), and the size or volume of the arena in which experimental trials were performed (for details see (Uiterwaal & DeLong 2020; Uiterwaal *et al*. n.d.)).

To obtain estimates of typical prey field densities, we used mass-abundance scaling relationships. Mass-abundance scaling relationships describe the general pattern that a species’ abundance is inversely related to its mass (Damuth 1981; White *et al*. 2007). Using the data from Hatton et al. (2019), we fit separate log-log regressions of abundance on body mass for mammals, birds, ectotherms, protists, and prokaryotes/algae in a Bayesian framework in Stan through the R package ‘*brms*’ using default priors. We then used the posterior predictive distributions of these models to estimate abundances for each prey species in the FoRAGE database whose body mass was available. Specifically, we determined each prey’s density at every decile of its posterior predictive distribution (the 10^th^ percentile to 90^th^ percentile by 10’s), and we used these abundances in our calculation of saturation in order to assess the sensitivity of the saturation index to the potential mis-estimation of prey abundances (see Supplemental Information S2 for details).

Prior to calculating the saturation index, we removed all studies from FoRAGE that used non-living prey or fungi as prey and studies without associated prey masses. During our analysis, we identified a small number of functional responses with prey abundance unit conversion issues in the data underlying FoRAGE. We then systematically checked all of the prey abundance unit conversions and removed those studies with incorrect conversions (74 functional responses from 20 studies; these will be corrected in future versions of FoRAGE). This reduced the original dataset from 2,598 functional responses to 2,100 which we refer to as the ‘full dataset’.

### Relationships between the Saturation Index and Covariates

To examine the relationships between the saturation index and potential covariates, we used generalized linear mixed effects models. As fixed effects, we included the major prey and predator taxa (phylum to class), habitat (terrestrial, aquatic-freshwater, and aquatic-marine), interaction dimension (2D, 2.5D, and 3D), arena size used in the experiment, the natural logarithm of prey mass, the natural logarithm of predator mass, and a quadratic effect of temperature. We included arena size because prior studies, including those using FoRAGE, have shown an effect of arena size on space clearance rates (Uiterwaal & DeLong 2018, 2020). We modeled temperature as a quadratic effect to allow for unimodal relationships with temperature (Englund *et al*. 2011; Uiterwaal & DeLong 2020). Although the saturation index includes the prey density that we estimated from prey mass, we included prey mass in our model because it is also influences space clearance rate and handling time and could therefore influence the saturation index through these parameters as well (Vucic-Pestic *et al*. 2010; Rall *et al*. 2012; Uiterwaal & DeLong 2020). To account for the non-independence of functional response estimates due to multiple estimates occurring on taxonomically similar species, we included minor prey and predator taxa (class to family) as random effects (a table of major and minor taxa is in Supplemental Information S3). Because the saturation index is limited to values between zero and one, we modeled the response as Beta distributed with a logit link function. We fit the regression model to the data in Stan through R using the package ‘*brms*’ with default priors.

We fit the model to a subset of the full dataset. We excluded studies that were missing any values for the covariates included in the analysis and those for which the fitted handling time value was less than 1×10^−6^ days to exclude functional responses with unidentifiable handling times (Uiterwaal & DeLong 2020). We also limited the number of major predator and prey taxa considered by dropping all predator taxa with less than 15 functional response studies and then dropping all prey taxa with less than 15 functional response studies. We limited the number of major predator and prey taxa to ensure enough functional response studies within each major taxa to include minor taxa as a random effect and prevent estimating major taxa effects based on only a few observations. Last, we also excluded mammals and birds for several reasons. First, the functional response studies on mammals and birds were performed in the field whereas all remaining studies were performed in laboratory conditions, confounding the effects of these taxa with the effects of measuring functional responses in the field. Second, the mammal and bird studies had no area boundaries, preventing our inclusion of arena size in the analysis. Third, unlike for all other taxa, which are ectotherms, the bird and mammal temperatures listed in FoRAGE represent the predator’s endothermic average body temperature rather than the environmental temperature at which the study was conducted. After applying these criteria to the 2,100 functional response studies in the full dataset, 1,468 studies remained. We used this ‘reduced dataset’ in our generalized linear mixed model analysis. We performed the analysis using the saturation index calculated at the 10^th^ percentile, median, and 90^th^ percentile of estimated prey densities.

### Explaining Relationships between the Saturation Index and Covariates

The analysis of the relationship between the saturation index and covariates gives the net relationship between saturation and the covariates but does not explain *why* these covariates have the relationships with saturation they do. As the saturation index is a function of the functional response parameters (space clearance rate and handling time) and the density of prey (Equation 3), the effects of the covariates on the saturation index are dependent on their effects on these three factors. We therefore performed two additional analyses on the reduced dataset using the same model as for estimating the effects of covariates on the saturation index to determine the partial effects of each of the covariates on the natural log of the space clearance rates and handling times. We again used Stan through R using the package ‘*brms*’ with default priors assuming a normal distribution of residuals and an identity link. We did not perform an analysis for prey densities as the response variable because these were determined completely by prey mass and classification as algae, ectotherm, or protist (see Methods: Estimating the Saturation Index).

## Results

### Estimates of Functional Response Saturation

The feeding rate saturation index showed a right-skewed distribution with a mode near zero for most deciles of prey densities (10^th^ through 70^th^ percentiles; Figure 2). At the two highest deciles of prey densities (80^th^ and 90^th^ percentiles; Figure 2), the saturation index showed a bimodal distribution with modes near zero and one. For the full dataset, half of the studies were below a saturation index of 0.002 at the 10^th^ percentile of prey densities, below 0.05 at the 50^th^ percentile, and below 0.57 at the 90^th^ percentile (Figure 2A). For the reduced dataset, half of the studies were below a saturation index value of 0.003 at the 10^th^ percentile, below 0.06 at the 50^th^ percentile, and below 0.62 at the 90^th^ percentile (Figure 2B-2E).

**Figure 2.**
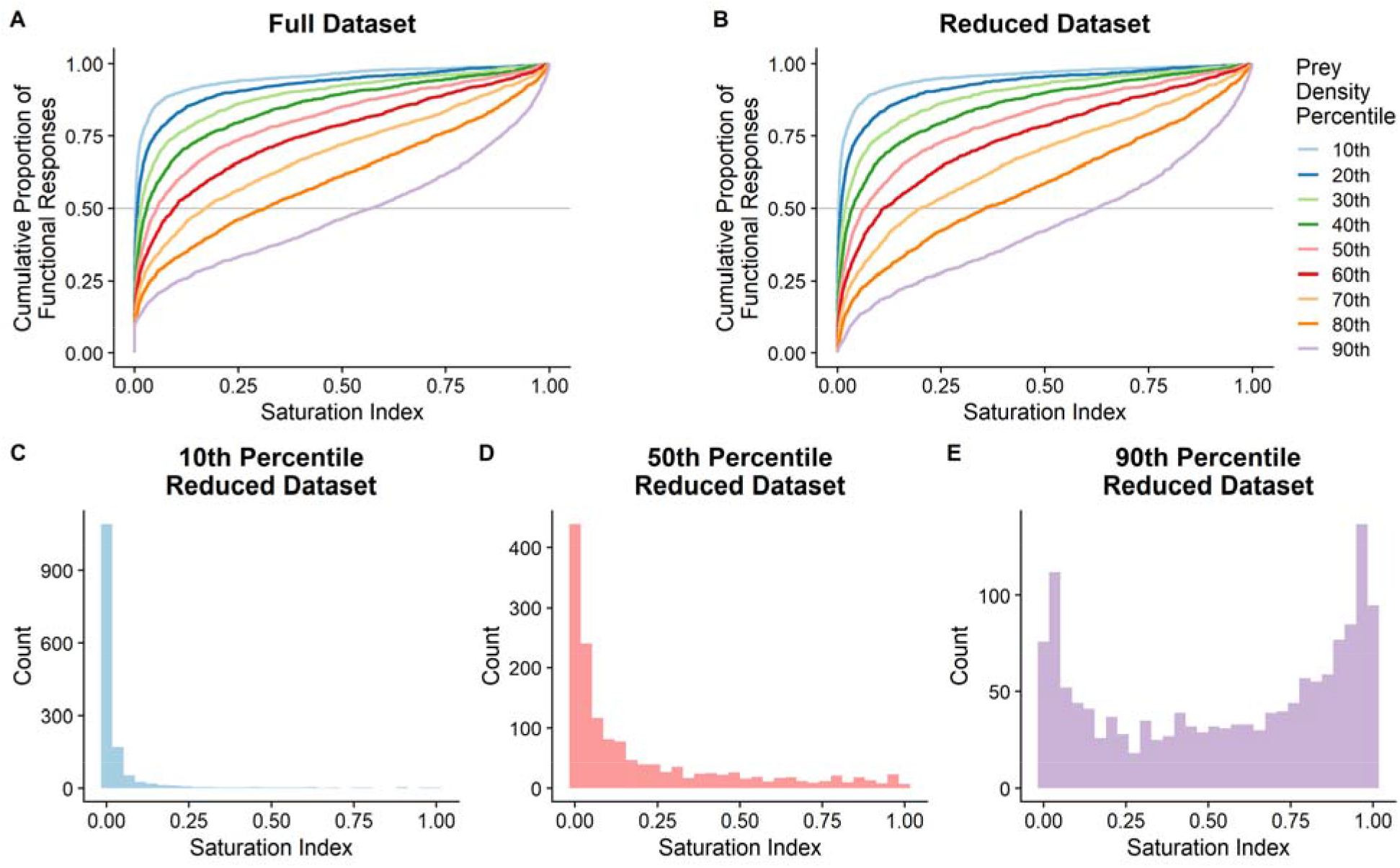
For both the full (A) and reduced (B) datasets, the index of predator feeding rate saturation showed right-skewed distributions with a mode near zero for the 10^th^ to 70^th^ percentiles of mass-estimated prey densities, and bimodal distributions with modes near zero and one for the 80^th^ and 90^th^ percentiles of prey densities. Histograms of the index distributions at the 10^th^, 50^th^, and 90^th^ percentiles of prey densities for the reduced dataset are given in C, D, and E, respectively.

### Analysis of Functional Response Saturation Covariates

Here we present only the results for the saturation index calculated at the median estimate of prey density. Similar results occurred at the 10^th^ and 90^th^ percentiles of estimated prey densities except for estimates of prey taxa partial effects (Supplementary Information S4 and Discussion). Our model, the intercept of which represents an amphibian feeding on algae in freshwater in three dimensions (−6.07; 90% Credible Interval (CrI) (−7.36,-4.82)), suggests that all considered covariates influence the feeding rate saturation (Figure 3; for a summary table of the regression results see Supplementary Information S3). As prey, amphibians (median posterior partial effect = 2.03; (0.95,3.1)), crustaceans (0.87; (0.22, 1.57)), fish (2.04; (1.38,2.78)), insects (1.38; (0.74,2.09)), and mollusks (1.89;(0.83,3.01)) showed positive partial effects on saturation with all other prey taxa showing no apparent partial effects (Figure 3A). As predators, only fish showed a positive partial effect on the saturation index (1.14;(0.47,1.8)) with all other predator taxa showing no apparent partial effects (Figure 3B). Marine habitats showed a positive partial effect on the saturation index (0.49; (0.25,0.73)) as did the interaction dimension being 2 (1.24; (0.97,1.52)) or 2.5D (0.94; (1.66,1.22); Figures 3C, 3D). For the continuous factors, the saturation index decreased with prey mass (−0.26; (−0.29,0.34); Figures 3E, 3I), increased with predator mass (0.04; (0.01,0.07); Figures 3F, 3J), showed a unimodal, concave relationship with temperature (median posterior linear effect = 0.05; (0.07,0.09); median posterior quadratic effect = -0.0018; (−0.003,-0.0008); Figures 3G, 3K), and increased with arena size (0.08; (0.04,0.12); Figures 3H, 3L).

**Figure 3.**
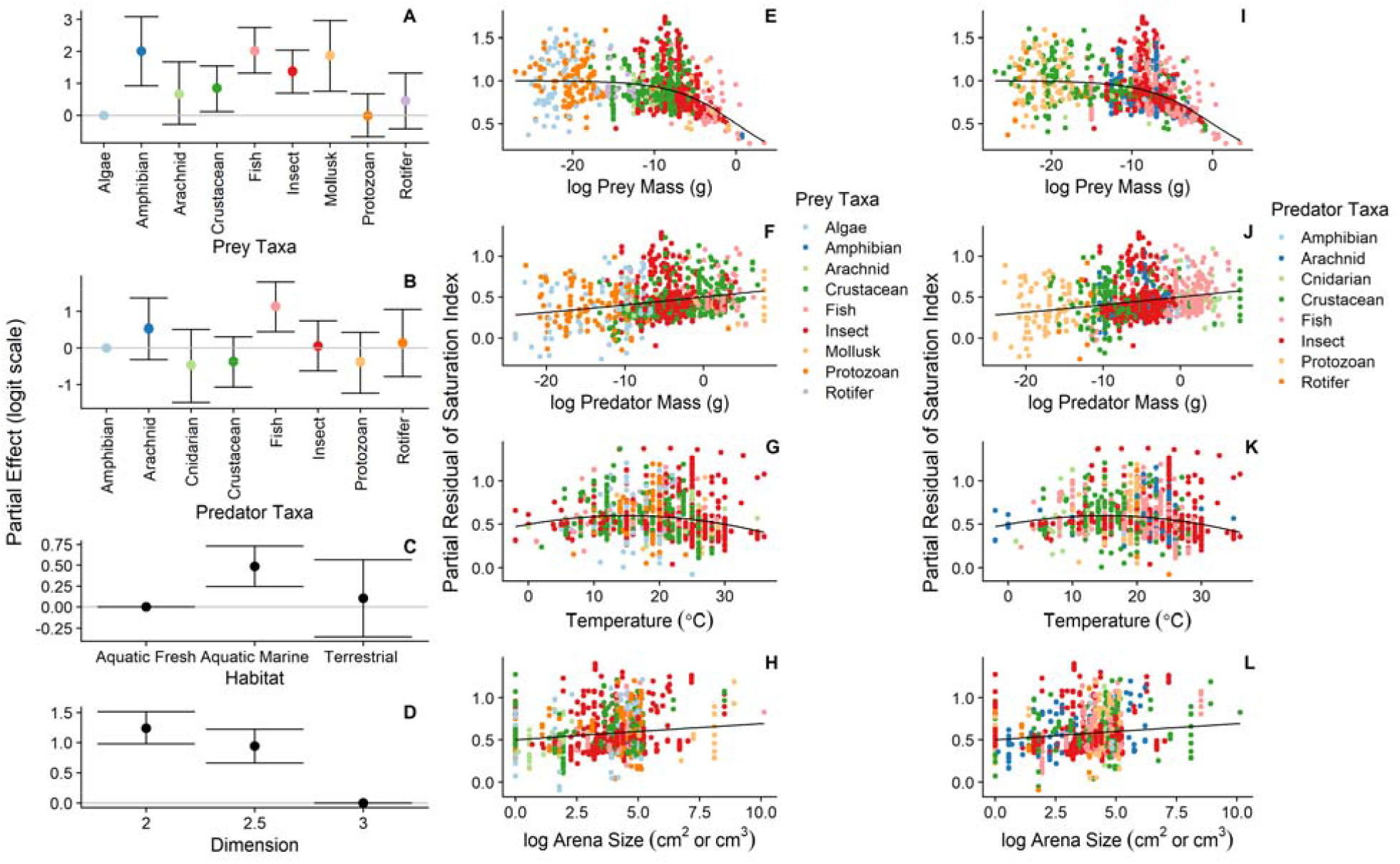
Prey taxa (A), predator taxa (B), habitat (C), and dimension (D) exhibited partial effects on the saturation index on the logit scale (error bars represent 90% credible intervals). The saturation index decreased with prey mass (E,I), increased with predator mass (F,J), showed a unimodal, concave relationship with temperature (G,K), and increased with arena size (H,L). Note that E-H and I-L include the same data, but E-H are color-coded by prey taxa and I-L are color-coded by predator taxa. Colors in E-H correspond to the same colors in A and the colors in I-L correspond to the same colors in B.

### Covariate Effects on Space Clearance Rates and Handling Times

With the intercept (−8.5; (−10.9,-6.2)) again representing an amphibian feeding on algae in freshwater in three dimensions, the model explaining variation in space clearance rates (Figure 4; See Supplemental Information S6 for a summary table of the regression results) suggested that arachnids (2.13; (0.02,4.29)), fish (2.15; (0.71,3.64)), insects (2.71, (1.24, 4.12)), and rotifers (2.37; (0.59, 4.09)) had partial positive effects on space clearance rates when they were the prey with all other prey showing no apparent partial effect(Figure 4A) and that fish had a partial positive effect on space clearance rate when they were predator (3.87, (1.86,5.9)) with all other predator taxa showing no apparent partial effect (Figure 4B). Marine habitats had a positive partial effect (0.78; (0.29,1.25)) while terrestrial habitats had a negative partial effect on space clearance rates (−2.71; (−3.63,-1.82); Figure 4C). Two- and 2.5-dimension studies showed positive partial effects on space clearance rates (2D: 4.61; (4.08,5.15), 2.5D: 3.99; (3.43,4.55); Figure 4D). Whereas predator mass was positively associated with space clearance rates and temperature showed a unimodal, concave relationship with space clearance rates, prey mass and arena size did not have statistically clear relationships with space clearance rates (Figures 4E-H).

**Figure 4.**
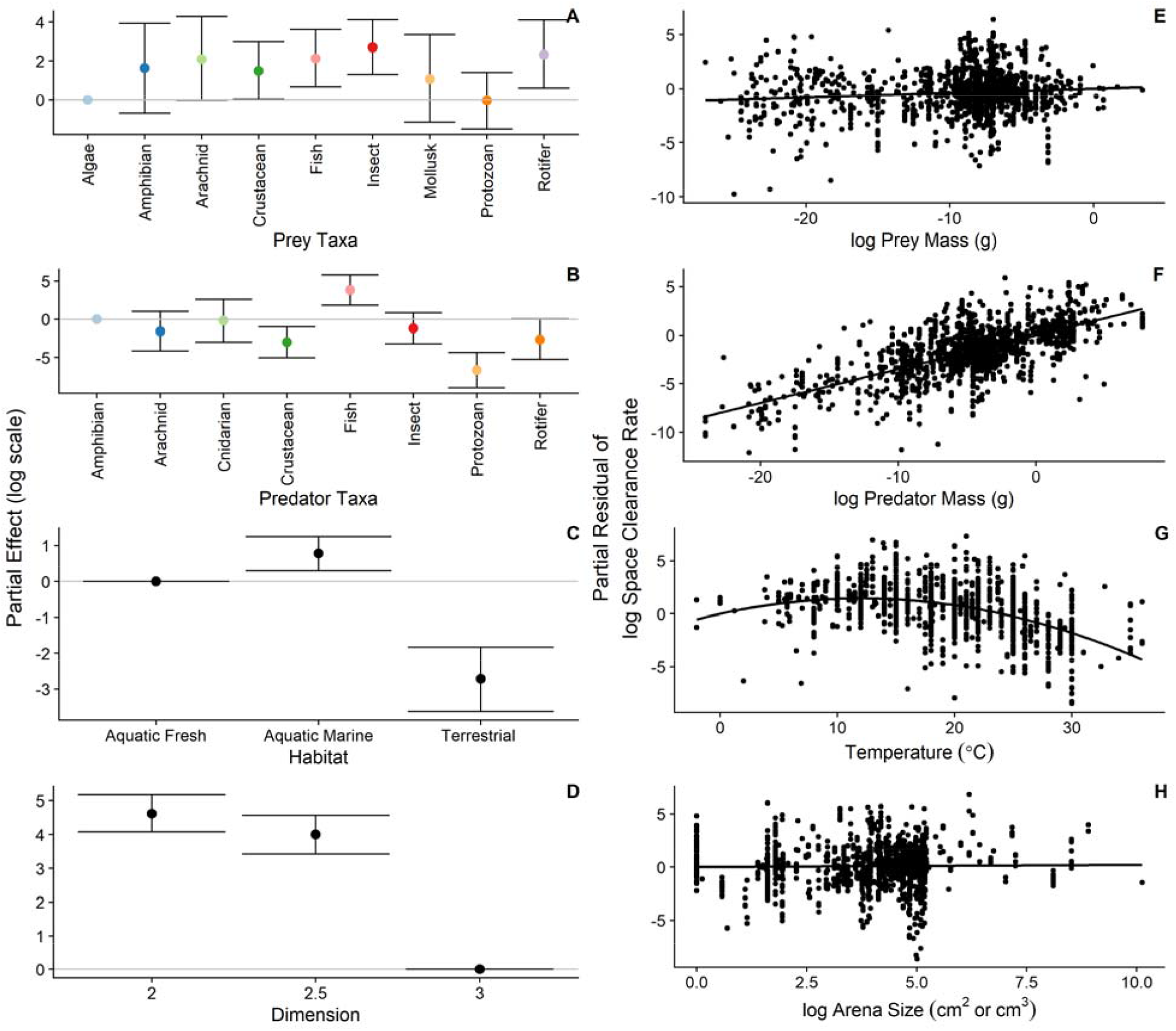
Prey taxa (A), predator taxa (B), habitat (C), and dimension (D) exhibited partial effects on log-transformed space clearance rates (error bars represent 90% credible intervals). Space clearance rates increased with predator mass (E), showed a unimodal, concave relationship with temperature (G), and had no apparent relationship with prey mass and arena size (D, H).

With the intercept (−10.9; (−13.0,-8.9)) still representing an amphibian feeding on algae in freshwater in three dimensions, the model explaining variation in handling times (Figure 5; See Supplemental Information S6 for a summary table of the regression results) suggested that all prey taxa other than algae showed positive partial effects on handling times (Amphibian: 9.3; (7.311.2), Arachnid: 6.4; (4.44,8.3), Crustacean: 7.0; (5.7,10.2), Fish: 8.93; (7.7,10.2), Insect: 6.7; (5.5,8.0), Mollusk: 9.8; (7.9,11.7), Protist: 2.26; (1.0,3.6); Rotifer: 5.98; (4.5,7.6); Figure 5A). As predators, fish showed a negative partial effect (−1.86; (−3.6,-0.12)) and protists showed a positive partial effect on handling times (6.0; (3.7,7.7)) with all other predator taxa showing no apparent partial effects (Figure 5B). Marine and terrestrial habitats showed positive partial effects (Marine: 0.4; (0.04,0.8), Terrestrial: 2.5; (1.8,3.2); Figure 5C) while 2- and 2.5-dimension interactions had negative partial effects on handling times (2D: - 1.1; (−1.5,-0.7), 2.5D: -1,3; (−1.7,-0.9); Figure 5D). For the continuous variables, handling times increased with prey mass, decreased with predator mass, increased with arena size, and decreased linearly with temperature (Figures 5E-H).

**Figure 5.**
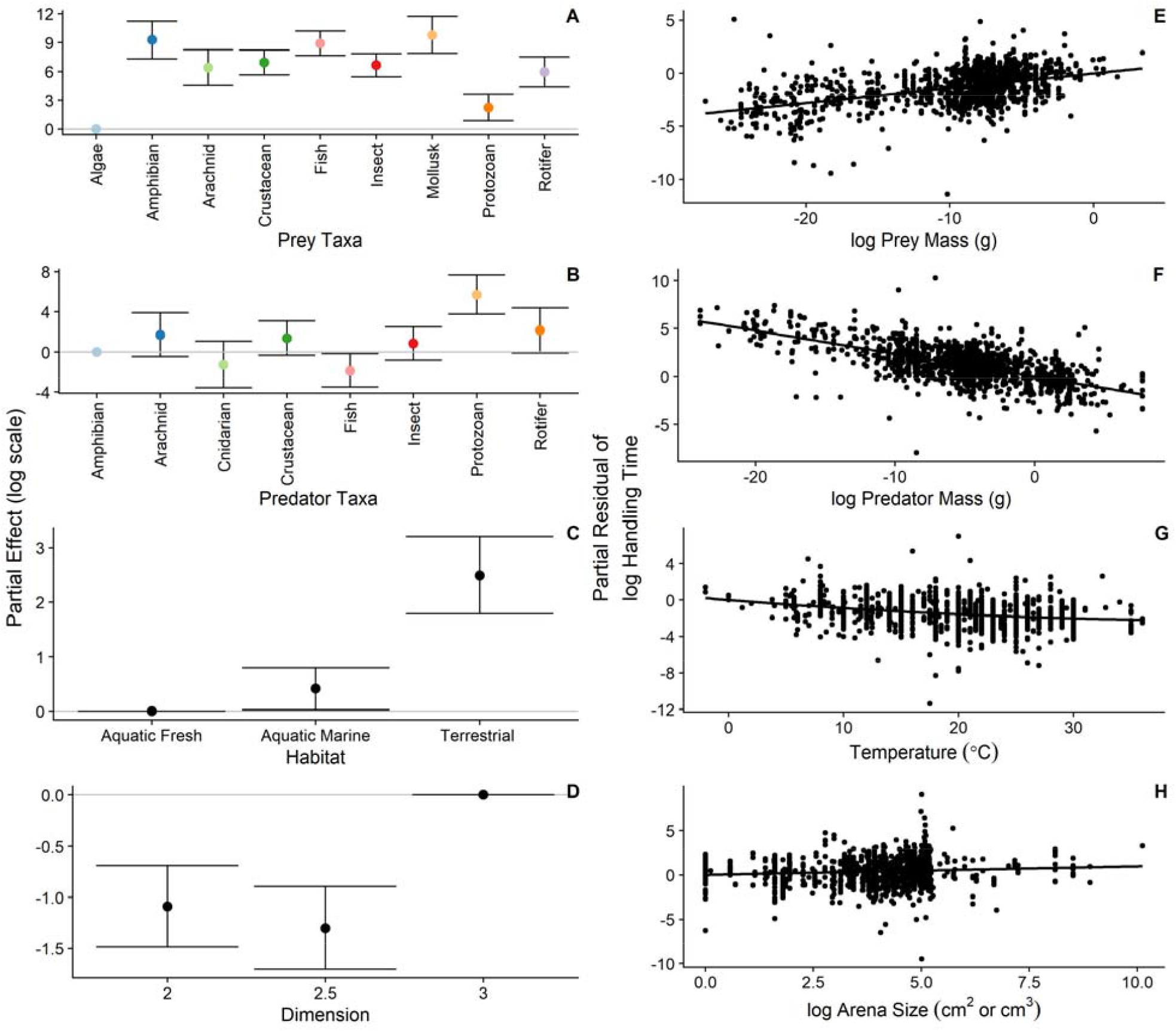
Prey taxa (A), predator taxa (B), habitat (C), and dimension (D) exhibited partial effects on log-transformed handling times (error bars represent 90% credible intervals). Handling times increased with prey mass (D), decreased with predator mass (E), decreased with temperature (G), and increased with arena size (H).

## Discussion

Combining functional response parameter estimates from laboratory-controlled experiments with field-relevant estimates of prey density obtained from mass-abundance scaling relationships, our results suggest that predator feeding rates may often be unsaturated under typical field conditions. Indeed, our analysis identified several predator functional responses that remained unsaturated even at high prey densities (i.e. 25% of the functional responses were below a saturation index value of 0.2 with prey densities at the 90^th^ percentile). For these functional responses, variation in feeding rates should be determined largely by variation in space clearance rates and prey densities. Space clearance rates are the product of predator and prey velocities, the distance over which predators can detect prey, the probability of a predator attacking a prey individual given its detection, and the probability that the attack is successful (Jeschke *et al*. 2002; DeLong 2021; Wootton *et al*. n.d.). For unsaturated feeding rates, these processes are central to determining the magnitude of feeding rates. A lack of saturation also means that predator feeding rates are lower than their potential maxima. This result is congruent with a previous study on fishes suggesting that fish digestive capacities are often larger than what would be necessary for the average amount of food they encounter (Armstrong & Schindler 2011). Feeding rates occurring below their potential maxima may also be indicative of the evolution of prudent predation (Gutiérrez Al-Khudhairy & Rossberg 2022) or constraints on predator feeding rates from other sources (Vuorinen *et al*. 2021). Last, unsaturated feeding rates should dynamically respond to changes in prey densities leading to density-dependent prey mortality that is close to proportional to prey densities. Our results suggest that for the species that remained unsaturated at the highest prey abundance decile, the use of linear functional responses to describe variation in feeding rates may be a sufficient approximation (Wootton & Emmerson 2005; Novak 2010; Jonsson 2017).

Many of the functional responses showed a gradient in feeding rate saturation with unsaturated feeding rates under most deciles of prey abundance and saturated feeding rates at the very highest deciles of prey abundance. In the full dataset, less than half of the functional responses show a reduction in feeding rates of more than 20% (relative their hypothetical linear functional response) when prey were assumed to be at their 70^th^ abundance percentile. Yet, with the prey at the 90^th^ abundance percentile, half of the functional responses have feeding rates that show a reduction of over 60%. This result suggests two nonexclusive scenarios for feeding rate saturation. One is that the extent of saturation may be dependent on whether a prey species is relatively abundant or not given its mass. For prey that are very abundant for their mass, predator feeding rates are likely to be saturated. Yet, for prey with abundances more typical of their size, predator feeding rates are likely to be unsaturated. For example, invasive and pest species can reach extremely high abundances relative to other species of their size (Hall Jr. *et al*. 2006). Predators feeding on these species may exhibit saturated feeding rates even though they might exhibit unsaturated feeding rates on prey of a similar size with more typical abundances. Another view of the result that feeding rate saturation occurs generally at the highest predicted prey abundances is that the predator’s feeding rate may be typically unsaturated but become saturated in times or areas where prey are particularly abundant. For example, extreme abundance events like oak masts and periodical cicada emergences are known to saturate predators and are thought to have evolved for that purpose (Karban 1982; Kelly 1994). Indeed, many field functional response studies show feeding rate saturation at only a few high prey abundance observations (Angerbjörn 1989; Korpimaki & Norrdahl 1991; Redpath & Thirgood 1999; Sundell *et al*. 2000; Quinn *et al*. 2017; Coblentz *et al*. n.d.). Thus, it may be that unsaturated predator feeding rates are typical except when prey exhibit high abundances.

We found that the extent of feeding rate saturation depended on prey and predator taxonomic identities and masses, habitat, interaction dimension, and temperature after accounting for experimental arena size. Our results suggest that, at the median prey abundances, amphibians, crustaceans, fish, insects, and mollusks showed a greater degree of saturation as prey than algae. These same five prey taxa also show positive partial effects on space clearance rates and handling times. However, our results suggest that the differences among prey taxa in there are effects on saturation are dependent on how abundant they are. For example, protists and rotifers showed negative partial effects at the 10^th^ percentile of estimated prey densities with all other taxa having no apparent effect and all prey taxa other than the reference taxa, algae, showed positive partial effects at the 90^th^ percentile of estimated prey densities (Supplemental Information S4). For the predator taxa, only fish showed a positive partial effect on the degree of saturation. This likely reflects the generally higher space clearance rates of fish relative to the other predator taxa, which has been attributed to their relatively higher velocities in moving through their environment after accounting for body size (Pawar *et al*. 2012; Buba *et al*. 2022; Wootton *et al*. n.d.). This conclusion is partially supported by a similarly positive partial effect for mammals and birds which also are likely to have higher velocities in their environments for their body sizes in an analysis including these predator taxa but not accounting for arena size (Supplementary Information S5).

Habitat and interaction dimension also had effects on the degree of feeding rate saturation, with 2- and 2.5-dimensional interactions having higher levels of feeding rate saturation compared to 3-dimensional interactions. This result is driven by the higher values of space clearance rates in the 2- and 2.5-dimensional studies that outweighed the generally lower handling times in 2- and 2.5-dimensional studies. In general, although the magnitudes of space clearance rates are not comparable across dimensions due to differences in spatial units (e.g. m^2^predator^-1^time^-1^ versus m^3^predator^-1^time^-1^, Uiterwaal & DeLong 2020), the saturation index is unitless. Therefore, the higher absolute values of space clearance rates in 2 and 2.5 dimensions lead to greater saturation. However, it remains unclear why 2- and 2.5-dimensional space clearance rates are generally greater than 3-dimensional space clearance rates. Marine studies also showed higher feeding rate saturation as a result of higher space clearance rates and handling times. Although terrestrial studies also showed higher space clearance rates compared to freshwater studies, they also showed lower handling times that counteracted the effects of higher space clearance rates. It is unclear whether this pattern of greater saturation in marine studies is general or whether it is the product of the species on which functional response studies have been conducted across different habitats. Confirming whether marine species show generally greater feeding rate saturation could be important for understanding how predation operates differently in different ecosystems (Shurin *et al*. 2002, 2006).

Overall, our results suggest that predator and prey masses are likely to have opposite net effects on the degree of feeding rate saturation. We found that prey mass was negatively associated with the degree of feeding rate saturation. However, previous research has shown that increasing prey mass is associated with higher space clearance rates and higher handling times, which would lead us to expect prey mass to be positively associated with feeding rate saturation (Vucic-Pestic *et al*. 2010; Rall *et al*. 2012; Uiterwaal & DeLong 2020). This difference in expectation can be explained by the fact that prey density decreases with prey mass (Damuth 1981; White *et al*. 2007; Hatton *et al*. 2019) and that the net effect of prey mass on feeding rate saturation is the product of all three of these relationships. That is, the negative scaling of prey densities with prey mass is stronger than the positive relationships between prey mass and space clearance rates and handling times, resulting in a positive effect of mass on saturation. In contrast to prey mass, predator mass was positively associated with feeding rate saturation. Previous results suggest that predator mass is typically positively associated with space clearance rates and negatively related to handling times (Vucic-Pestic *et al*. 2010; Rall *et al*. 2012; Uiterwaal & DeLong 2020). In our dataset, predator mass exhibits a slightly stronger positive relationship with space clearance rates than a negative relationship with handling times, thereby producing a net positive relationship between feeding rate saturation and predator mass.

Temperature has strong effects on functional response parameters in laboratory studies (Thompson 1978; Englund *et al*. 2011; Rall *et al*. 2012; Uiterwaal & DeLong 2020), with studies typically documenting positive or unimodal, concave relationships between temperature and space clearance rates and negative or unimodal, convex relationships between temperature and handling times (Englund *et al*. 2011; Rall *et al*. 2012; Uiterwaal & DeLong 2020). This suggests that the net effect of temperature on feeding rate saturation should be dependent on the relative strengths of the relationships between temperature and space clearance rates and handling times. Our results show a stronger unimodal, concave relationship between space clearance rates and temperature than the negative relationship between handling times and temperature, and this leads to a net unimodal, concave relationship between temperature and saturation. These results lead to the prediction of a mid-latitudinal peak in feeding rate saturation and that the degree of feeding rate saturation will be sensitive to continued climate change, with potentially profound consequences for predator-prey interactions on a global scale.

Although our results suggest that predator feeding rates are unsaturated across a range of typical prey densities, many of our estimates are likely to be overestimates. First, the mass-abundance scaling relationships we used to predict prey densities may overestimate prey densities because the abundances are typically reported in aerial square meters. Thus, for aquatic organisms, abundances are integrated over some depth often greater than a meter and therefore are likely overestimates of the abundances in a cubic meter, the relevant metric for three dimensional studies (Hatton *et al*. 2019). Mass-abundance scaling relationships also may reflect maximum abundances rather than typical abundances because researchers often measure abundances of organisms where they are abundant (Lawton *et al*. 1990; Marquet *et al*. 1995; White *et al*. 2007). Second, because functional response experiments are typically performed in spatially and structurally simplified arenas, the estimates of functional response parameters may be biased toward values that lead to higher feeding rates than those that are likely to be observed in nature (Novak *et al*. 2017; Griffen 2021).

In our analysis, we assumed that predator feeding rates within predator-prey pairs were described by a saturating Type II functional response. However, predators can exhibit other functional response types and typically incorporate more than one prey type into their diets. For example, predators might exhibit sigmoidal Type III functional responses or predator feeding rates could be dependent on predator densities (Holling 1959; DeLong & Vasseur 2011; Novak & Stouffer 2021). In general, considering these additional aspects of predator functional responses shows that our estimates of saturation will be conservative or show little change with these alternative functional response scenarios (See Supplemental Information S7 for a general derivation of the saturation index and specific examples). In the case of a Type III functional response, the saturation index becomes a sigmoidal function of prey densities and should give similar results as the Type II functional response except with lower saturation values at low prey densities (Supplemental Information S7). In the case of functional responses with predator dependence or the inclusion of multiple prey in the predator’s diet, the degree of feeding rate saturation should be lower than that estimated for the Type II functional response and our results here will be conservative (Supplemental Information S7). However, one caveat with respect to the multi-prey case is that, although feeding rate saturation with respect to the focal prey should decrease with the addition of alternative prey, the saturation of the predator’s total feeding rate across all prey can increase with the addition of alternative prey. Whether saturation of the predator’s total feeding rate increases with additional prey in the diet will depend on whether and how the parameters of the functional response change with the addition of prey species to the diet. In general, we know little about how functional response parameters are likely to change with diet richness and understanding how total feeding rate saturation in the predator is likely to change with diet richness will require studies measuring functional responses and their saturation under field conditions.

## Conclusion

The degree to which predator feeding rates are saturated has important consequences for what factors predominantly determine predator feeding rates, whether predator feeding rates are near their maxima or not, and how predator-prey interaction strengths respond to changes in prey densities. Our results suggest that it may be the case that predator feeding rates are often far from saturated over large ranges of typical prey densities. Furthermore, our results suggest that the degree of feeding rate saturation is shaped by predator and prey traits and the environment. We suggest that future work on feeding rate saturation focus on 1) measuring saturation under field conditions, 2) understanding the proximate and ultimate causes of feeding rates being unsaturated over a range of typical prey densities, and 3) determining the ecological and evolutionary consequences of unsaturated feeding rates for predator-prey systems.

## Acknowledgements

This work was supported by a James S. McDonnell Foundation Scholar Award in Studying Complex Systems to JPD.

