## Supplemental Information for "Predator feeding rates may often be unsaturated under typical prey densities"

**Contents** **Page**


| | |
|---|---|
| **S1 Deriving the Index of Saturation** | **2** |
| **S2 Mass-Abundance Scaling** | **3** |
| **S3 Analysis of Saturation Index Covariates** | **6** |
| **S4 Analysis Results of Saturation Index Covariates for High and Low Prey Densities** | **9** |
| **S5 Analysis Results of Model Including Birds and Mammals** | **13** |
| **S6 Analysis of Space Clearance Rate and Handling Time Covariates** | **15** |
| **S7 Generalization of Results to Common Functional Response Forms** | **17** |
| **Literature Cited** | **21** |

### S1 Deriving the Index of Saturation

The index of functional response saturation that we use ($I$) is the proportional reduction in predator feeding rate between the case that the predator would exhibit a linear Type I functional response ($f_I$; no saturation with respect prey density) and the case that the predator exhibits a saturating Type II functional response ($f_{\mathrm{II}}$; feeding rates saturate with the prey density). We express it as

$$I = \frac{f_I - f_{II}}{f_I} \qquad \text{eqn. S1.1}$$

where

$$f_I = a\mathrm{R}, \qquad \text{eqn. S1.2}$$

$$f_{\mathrm{II}} = \frac{a\mathrm{R}}{1+ah\mathrm{R}}, \qquad \text{eqn. S1.3}$$

$a$ is the space clearance or attack rate, $h$ is the handling time, and R is the density of the prey. To see how we arrive at the formula for $I$ given in the main text eqn. 3, we provide the algebraic steps below beginning with the equation derived from substituting equations S1.2 and S1.3 into equation S1.1,

$$I = \frac{a\mathrm{R} - \frac{a\mathrm{R}}{1+ah\mathrm{R}}}{aR} \qquad \text{eqn. S1.4}$$

$$= 1 - \frac{1}{1+ah\mathrm{R}}\,. \qquad \text{eqn. S1.5}$$

We then replace the 1 in the first term with $\frac{1+ah\mathrm{R}}{1+ah\mathrm{R}}$ to get

$$= \frac{1+ah\mathrm{R}}{1+ah\mathrm{R}} - \frac{1}{1+ah\mathrm{R}}\,. \qquad \text{eqn. S1.6}$$

Because the two terms have the same denominator, we can bring them together to give

$$= \frac{ah\mathrm{R}}{1+ah\mathrm{R}}\,. \qquad \text{eqn. S1.7}$$

This is the same formula for $I$ given in the main text eqn. 3.

### S2 Mass-Abundance Scaling

Hatton et al. (2019) compiled a database of 5,985 observations of species abundances in units of aerial densities (number $m^{-2}$) and masses (g) across eukaryotes for the purpose of examining mass-abundance scaling relationships within and among taxa. Mass-abundance scaling shows a general negative relationship between the log abundance of organisms and their log mass. As the degree to which a functional response is saturated depends on the density of prey, we used the mass-abundance scaling relationships from the Hatton et al. dataset to estimate abundances for the prey species that occur in FoRAGE based on their masses. Rather than use the scaling relationships already calculated by Hatton et al. (2019) in their work, we refit the abundance-mass scaling relationships in a Bayesian framework. We did this so that we could use the posterior predictive distributions of the models to generate decile estimates of prey abundances and thereby assess the sensitivity of our inferences of feeding rate saturation. Posterior predictive distributions provide predicted future observations from a model for a set of observed predictor variables while incorporating both uncertainty in the parameter estimates of the model and the residual variance of the model (analogous to using prediction rather than confidence intervals in frequentist statistics).

We fit separate regressions of log abundance on log mass in grams for mammals (number of observations (n) = 2,852), birds (n = 603), ectotherms (n = 1,182), protists (n = 301), and algae/prokaryotes (n = 635). (Note that all of the algae/prokaryote observations come from a single study (Li 2002) and that the abundances actually represent the abundance of all organisms less than 20 µm in diameter, including both bacteria and algae). Graphs of each of the fitted scaling relationships along with the posterior predictive intervals used to generate the estimates of prey abundances are shown in Figures S2.1 and the regression results are provided in Table S2.1.

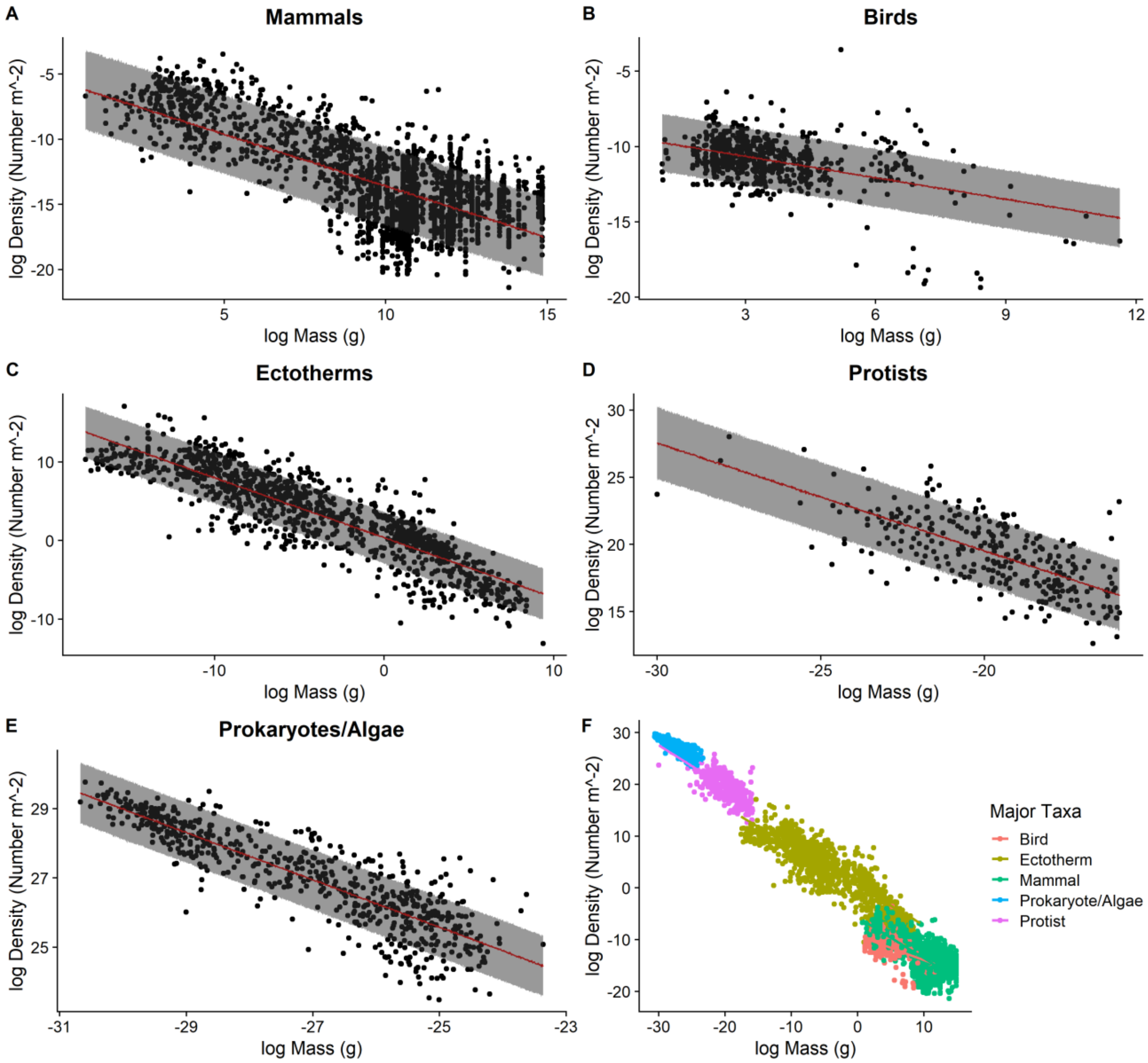


**Figure S2.1.** Mass-abundance scalings give estimated densities as a function of mass for mammals (A), Birds (B), Ectotherms (C), Protists (D), and Prokaryotes/Algae (E), and across all taxa (F). The red lines in A-E represent the median predicted densities while the shaded areas represent 80% prediction intervals (i.e. the upper and lower bounds are the predicted densities at the $90^{th}$ and $10^{th}$ percentile).

**Table S2.1.** The estimated regression parameters, 90% Credible Intervals (CrI) and Bayesian $R^2$ values of the mass-abundance scaling for each major taxon.

| Taxa | Intercept | 90% CrI | log Mass Coefficient | 90% CrI | Residual Standard Deviation | 90% CrI | Bayesian $R^2$ |
|---|---|---|---|---|---|---|---|
| Mammals | -5.66 | (-5.93,-5.37) | -0.79 | (-0.82,-0.77) | 2.37 | (2.31,2.42) | 0.48 |
| Birds | -9.23 | (-9.49,-8.97) | -0.47 | (-0.54,-0.41) | 1.48 | (1.41,1.56) | 0.19 |
| Ectotherms | 0.36 | (0.23,0.5) | -0.76 | (-0.78,-0.74) | 2.55 | (2.46,2.63) | 0.79 |
| Protists | 3.5 | (1.95,5.09) | -0.8 | (-0.88,-0.72) | 2.02 | (1.89,2.16) | 0.47 |
| Prokaryotes/ Algae | 8.54 | (7.86,9.23) | -0.68 | (-0.71,-0.66) | 0.68 | (0.65,0.71) | 0.77 |

### S3 Analysis of Saturation Index Covariates

Here we give the details of the model used to examine the relationship between covariates and the saturation index. Table S.3.1 gives the major and minor predator and prey taxa used in the analysis along with their associated sample sizes. Table S3.2 summarizes the regression coefficient estimates of the model.

**Table S3.1.** The major taxa used as fixed effects and minor taxa used as random effects in the analysis of the relationship between covariates and the saturation index. The number within parentheses after the taxa is the number of functional response studies on that taxa within the dataset.

| Predator Taxa | Prey Taxa |
|---|---|
| Amphibian (24) | Algae (81) |
| Anura (12) | Chlorophyte (26) |
| Urodela (12) | Cryptophyte (17) |
| Arachnid (122) | Diatom (20) |
| Mite (50) | Haptophyte (13) |
| Spider (72) | Ochrophyte (5) |
| Cnidarian (28) | Amphibian (10) |
| Hydrozoa (6) | Anura (4) |
| Scyphozoa (22) | Urodela (6) |
| Crustacean (310) | Arachnid (56) |
| Amphipod (66) | Mite (52) |
| Branchiopod (4) | Spider (4) |
| Cladoceran (27) | Crustacean (505) |
| Copepod (149) | Amphipod (55) |
| Decapod (27) | Branchiopoda (36) |
| Isopod (15) | Cladoceran (295) |
| Mysid (20) | Copepod (87) |
| Ostracod (2) | Decapod (2) |
| Fish (271) | Isopod (28) |
| Beloniformes (1) | Mysid (2) |
| Clupeiformes (15) | Fish (76) |
| Cypriniformes (34) | Mixed Taxa (2) |
| Cyprinodontiformes (10) | Atherinopsidae (2) |
| Gadiformes (2) | Centrarchidae (7) |
| Gasterosteiformes (5) | Cichlidae (6) |
| Perciformes (145) | Clupeidea (6) |
| Pleuronectiformes (7) | Cyprinidae (5) |
| Salmoniformes (38) | Gadidae (2) |
| Scorpaeniformes (4) | Moronidae (2) |
| Siluriformes (1) | Osmeridae (15) |
| Insect (617) | Perciformes (20) |
| Coleoptera (205) | Pleuronectidae (3) |
| Dermaptera (4) | Pleuronectiformes (1) |
| Diptera (69) | Poeciliidae (4) |
| Hemiptera (208) | Sciaenidae (1) |
| Heteroptera (18) | Insect (583) |
| Hymenoptera (10) | Chaoboridae (17) |
| Neuroptera (6) | Coenagrionidae (11) |
| Insects cont'd | Insects cont'd |

| | |
|---|---|
| Odonata (88) | Coleoptera (48) |
| Thysanoptera (5) | Culicidae (23) |
| Trichoptera (2) | Diptera (142) |
| Protozoan (84) | Ephemeroptera (9) |
| Chrysophyte (4) | Hemiptera (225) |
| Ciliate (37) | Heteroptera (4) |
| Dinoflagellate (38) | Homoptera (6) |
| Kateblepharidae (2) | Hymenoptera (5) |
| Sarcodine (3) | Lepidoptera (44) |
| Rotifer (23) | Odonata (3) |
| Asplanchnidae (11) | Orthoptera (5) |
| Brachionidae (12) | Thysanoptera (41) |
| | Mollusk (19) |
| | Bivalve (15) |
| | Gastropod (4) |
| | Protozoan (106) |
| | Chrysophyte (3) |
| | Ciliate (26) |
| | Dinoflagellate (72) |
| | Euglenid (2) |
| | Heterokont (3) |
| | Rotifer (32) |
| | Asplanchnidae (1) |
| | Brachionidae (20) |
| | Floscularíaceae (5) |
| | Synchaetidae (6) |

**Table S3.2**. The estimated parameters and 90% credible intervals (CrI) for each term in the model of the effects of covariates on the index of feeding rate saturation.

| Model Term | Estimate | 90% CrI |
|---|---|---|
| **Fixed Effects** | | |
| Intercept<br>(Corresponds to an Amphibian feeding on Algae in freshwater in three dimensions) | -6.07 | (-7.36,-4.82) |
| *Prey Major Taxa Effects* | | |
| Amphibian | 2.03 | (0.95, 3.1) |
| Arachnid | 0.66 | (-0.34, 1.67) |
| Crustacean | 0.87 | (0.22, 1.57) |
| Fish | 2.04 | (1.38, 2.78) |
| Insect | 1.38 | (0.74, 2.09) |
| Mollusk | 1.89 | (0.83, 3.01) |
| Protist | -0.01 | (-0.66, 0.67) |
| Rotifer | 0.45 | (-0.33, 1.3) |
| *Predator Major Taxa Effects* | | |
| Arachnid | 0.53 | (-0.28, 1.35) |
| Cnidarian | -0.46 | (-1.45, 0.52) |
| Crustacean | -0.39 | (-1.09, 0.29) |
| Fish | 1.14 | (0.47, 1.8) |
| Insect | 0.05 | (-0.62, 0.72) |
| Protist | -0.39 | (-1.23, 0.41) |
| Rotifer | 0.13 | (-0.8, 1.06) |
| *Habitat Effects* | | |
| Aquatic-Marine | 0.49 | (0.25, 0.73) |
| Terrestrial | 0.11 | (-0.35, 0.57) |
| *Dimension Effects* | | |
| 2D | 1.24 | (0.97, 1.52) |
| 2.5D | 0.94 | (0.66, 1.22) |
| *Continuous Variables* | | |
| log Prey Mass (g) | -0.26 | (-0.29, -0.23) |
| log Predator Mass (g) | 0.04 | (0.01, 0.07) |
| Temperature (C) | 0.05 | (0.01, 0.09) |
| $Temperature^2$ | -0.0018 | (-0.0028, -0.0008) |
| Arena Size | 0.08 | (0.04, 0.12) |
| **Random Effects** | | |
| Prey Minor Taxa Standard Deviation | 0.53 | (0.38, 0.70) |
| Predator Minor Taxa Standard Deviation | 0.40 | (0.26, 0.58) |
| **Distribution Parameters** | | |
| Beta Distribution Phi | 3.06 | (2.87, 3.27) |

#### S4 Analysis Results of Saturation Index Covariates for High and Low Prey Densities

In the main text, we only show the results of the analysis of the effects of covariates on the index of saturation for the median prey densities. Here we give the results for the low and high prey density deciles and show that the overall results are qualitatively similar to the analysis with the median prey densities except for some differences in whether the credible intervals for some predator and prey taxa overlap zero. Figs. S4.1 and S4.2 are equivalent to Figure 2 of the main text and Tables S4.1 and S4.2 are equivalent to Table S3.2.

*Low Prey Density Predictions*

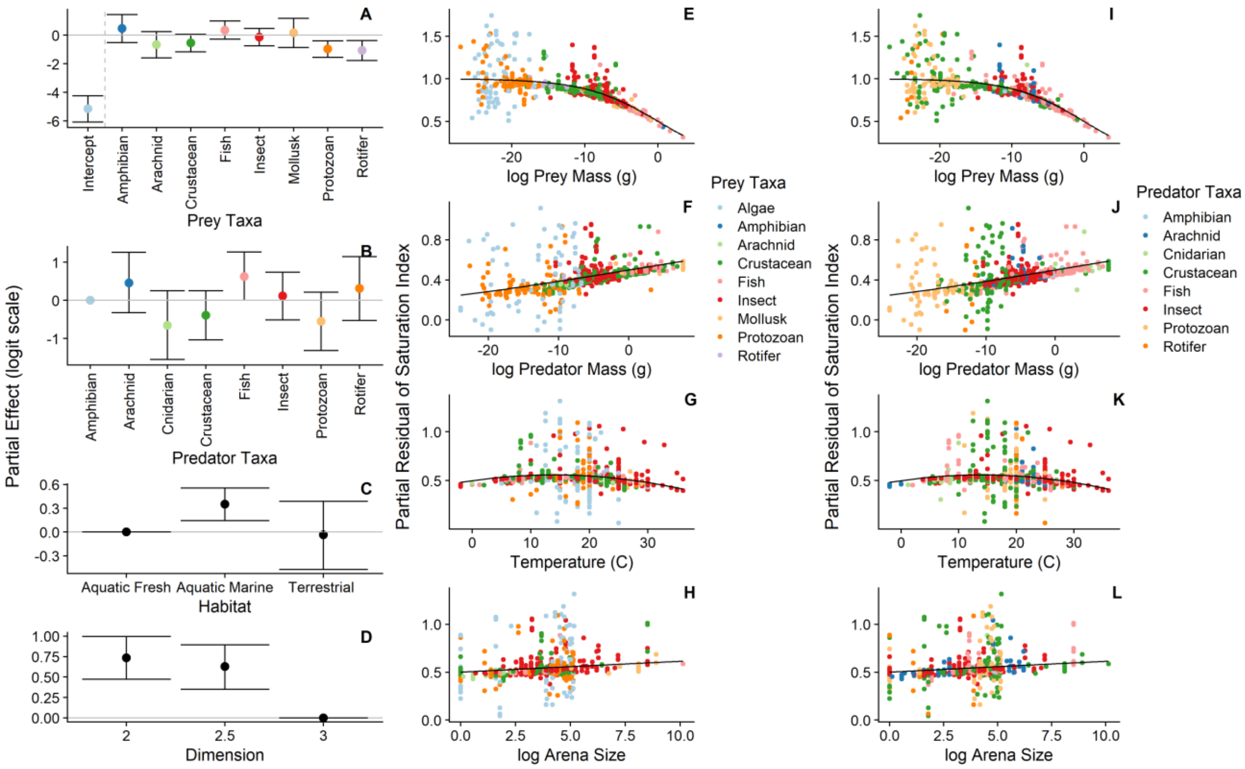


**Figure S4.1. For the low prey density predictions, prey taxa (A), predator taxa (B), habitat (C), and dimension (D) exhibited partial effects on the saturation index on the logit scale (the error bars represent 90% credible intervals). The saturation index decreased with prey mass (E,I), increased with predator mass (F,J), showed a unimodal, concave relationship with temperature (G,K), and an increasing relationship with arena size (H, L). Note that E-H and I-L include the same data, but E-H are color-coded by prey taxa and I-L are color-coded by predator taxa.**

**Table S4.1.** The estimated parameters and 90% credible intervals (CrI) for each term in the model of the effects of covariates on the index of saturation at the 10th percentile of predicted prey densities.

| **Model Term** | **Estimate** | **90% CrI** |
|---|---|---|
| **Fixed Effects** | | |
| Intercept<br>(Corresponds to an Amphibian feeding on Algae in freshwater in three dimensions) | -5.15 | (-6.08,-4.23) |
| *Prey Major Taxa Effects* | | |
| Amphibian | 0.48 | (-0.51, 1.43) |
| Arachnid | -0.67 | (-1.58, 0.25) |
| Crustacean | -0.55 | (-1.16, 0.06) |
| Fish | 0.35 | (-0.27, 0.99) |
| Insect | -0.14 | (-0.74, 0.46) |
| Mollusk | 0.16 | (-0.86, 1.17) |
| Protist | -0.97 | (-1.55, -0.41) |
| Rotifer | -1.06 | (-1.77, -0.37) |
| *Predator Major Taxa Effects* | | |
| Arachnid | 0.46 | (-0.32, 1.26) |
| Cnidarian | -0.65 | (-1.56, 0.25) |
| Crustacean | -0.39 | (-1.04, 0.25) |
| Fish | 0.62 | (0.001, 1.27) |
| Insect | 0.12 | (-0.51, 0.74) |
| Protist | -0.55 | (-1.32, 0.22) |
| Rotifer | 0.32 | (-0.53, 1.15) |
| *Habitat Effects* | | |
| Aquatic-Marine | 0.35 | (0.14, 0.55) |
| Terrestrial | -0.03 | (-0.47, 0.39) |
| *Dimension Effects* | | |
| 2D | 0.73 | (0.47, 1.00) |
| 2.5D | 0.63 | (0.35, 0.89) |
| *Continuous Variables* | | |
| log Prey Mass (g) | -0.20 | (-0.23, -0.18) |
| log Predator Mass (g) | 0.05 | (0.02, 0.07) |
| Temperature (C) | 0.03 | (-0.004, 0.07) |
| $\text{Temperature}^2$ | -0.001 | (-0.002, -0.0002) |
| Arena Size | 0.05 | (0.01, 0.08) |
| **Random Effects** | | |
| Prey Minor Taxa Standard Deviation | 0.45 | (0.34, 0.59) |
| Predator Minor Taxa Standard Deviation | 0.36 | (0.24, 0.51) |
| **Distribution Parameters** | | |
| Beta Distribution Phi | 6.77 | (6.16, 7.43) |

*High Prey Density Predictions*

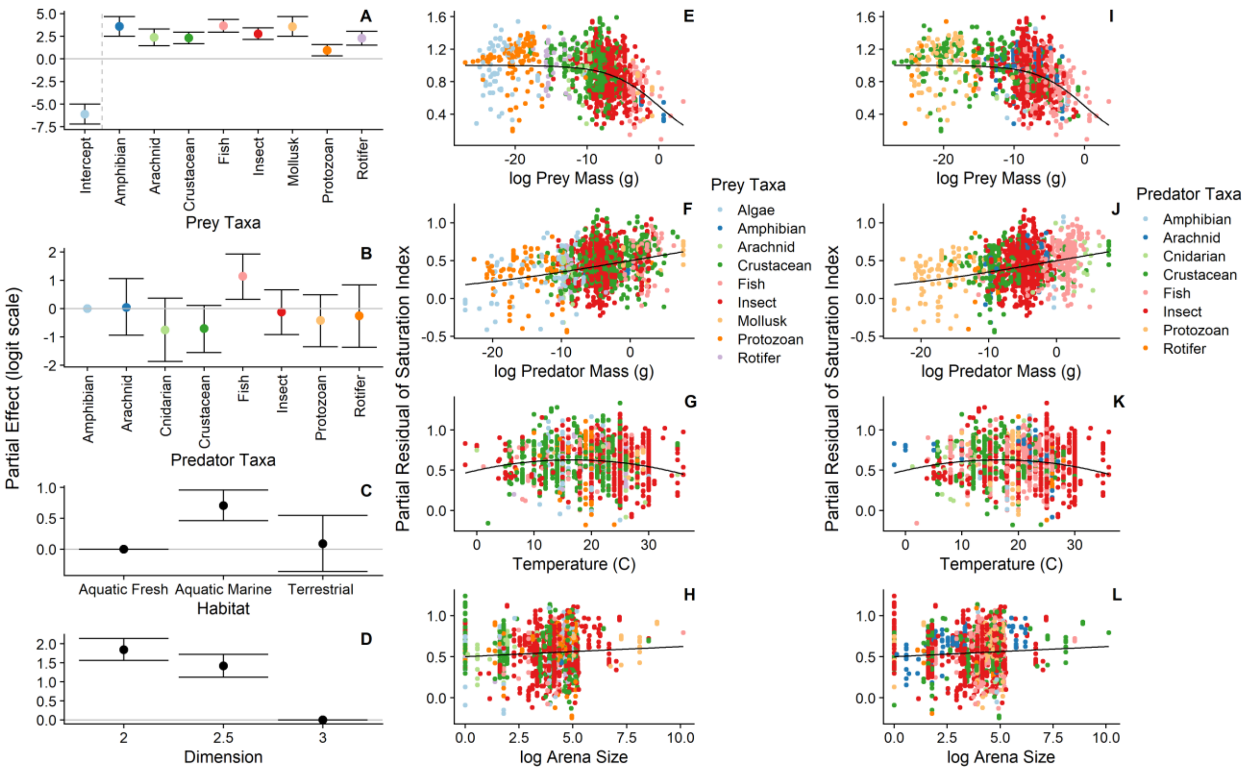


**Figure S4.2. For the high prey density predictions, prey taxa (A), predator taxa (B), habitat (C), and dimension (D) exhibited partial effects on the saturation index on the logit scale (the error bars represent 90% credible intervals). The saturation index decreased with prey mass (E,I), increased with predator mass (F,J), showed a unimodal, concave relationship with temperature (G,K), and showed a positive relationship with arena size (H,L). Note that E-H and I-L include the same data, but E-H are color-coded by prey taxa and I-L are color-coded by predator taxa.**

**Table S4.2.** The estimated parameters and 90% credible intervals (CrI) for each term in the model of the effects of covariates on the index of saturation at high predicted prey densities.

| **Model Term** | **Estimate** | **90% CrI** |
|---|---|---|
| **Fixed Effects** | | |
| Intercept (Corresponds to an Amphibian feeding on Algae in freshwater in three dimensions) | -6.1 | (-7.25, -4.98) |
| *Prey Major Taxa Effects* | | |
| Amphibian | 3.55 | (2.44, 4.64) |
| Arachnid | 2.37 | (1.43, 3.33) |
| Crustacean | 2.3 | (1.64, 2.99) |
| Fish | 3.61 | (2.88, 4.33) |
| Insect | 2.75 | (2.07, 3.43) |
| Mollusk | 3.53 | (2.45, 4.61) |
| Protist | 0.93 | (0.29, 1.58) |
| Rotifer | 2.26 | (1.48, 3.05) |
| *Predator Major Taxa Effects* | | |
| Arachnid | 0.06 | (-0.96, 1.05) |
| Cnidarian | -0.74 | (-1.93, 0.39) |
| Crustacean | -0.69 | (-1.57, 0.13) |
| Fish | 1.15 | (0.32, 1.95) |
| Insect | -0.11 | (-0.94, 0.69) |
| Protist | -0.40 | (-1.39, 0.55) |
| Rotifer | -0.24 | (-1.34, 0.81) |
| *Habitat Effects* | | |
| Aquatic-Marine | 0.71 | (0.45, 0.96) |
| Terrestrial | 0.09 | (-0.37, 0.53) |
| *Dimension Effects* | | |
| 2D | 1.84 | (1.55, 2.12) |
| 2.5D | 1.41 | (1.11, 1.70) |
| *Continuous Variables* | | |
| log Prey Mass (g) | -0.30 | (-0.33, -0.27) |
| log Predator Mass (g) | 0.06 | (0.04, 0.09) |
| Temperature (C) | 0.06 | (0.03, 0.1) |
| Temperature$^2$ | -0.002 | (-0.003, -0.001) |
| Arena Size | 0.05 | (0.01, 0.09) |
| **Random Effects** | | |
| Prey Minor Taxa Standard Deviation | 0.47 | (0.33, 0.63) |
| Predator Minor Taxa Standard Deviation | 0.49 | (0.33, 0.68) |
| **Distribution Parameters** | | |
| Beta Distribution Phi | 2.28 | (2.16, 2.42) |

### S5 Analysis Results of Model Including Birds and Mammals

In the main text, we analyze a reduced dataset that excluded birds and mammals. Here we give the results of the analysis with birds and mammals but without including arena size as a covariate (which is not applicable to birds and mammals as all studies considering them are field studies). Results show that birds and mammals (or the confounded effect of their field setting) have positive partial effects on the degree of feeding rate saturation.

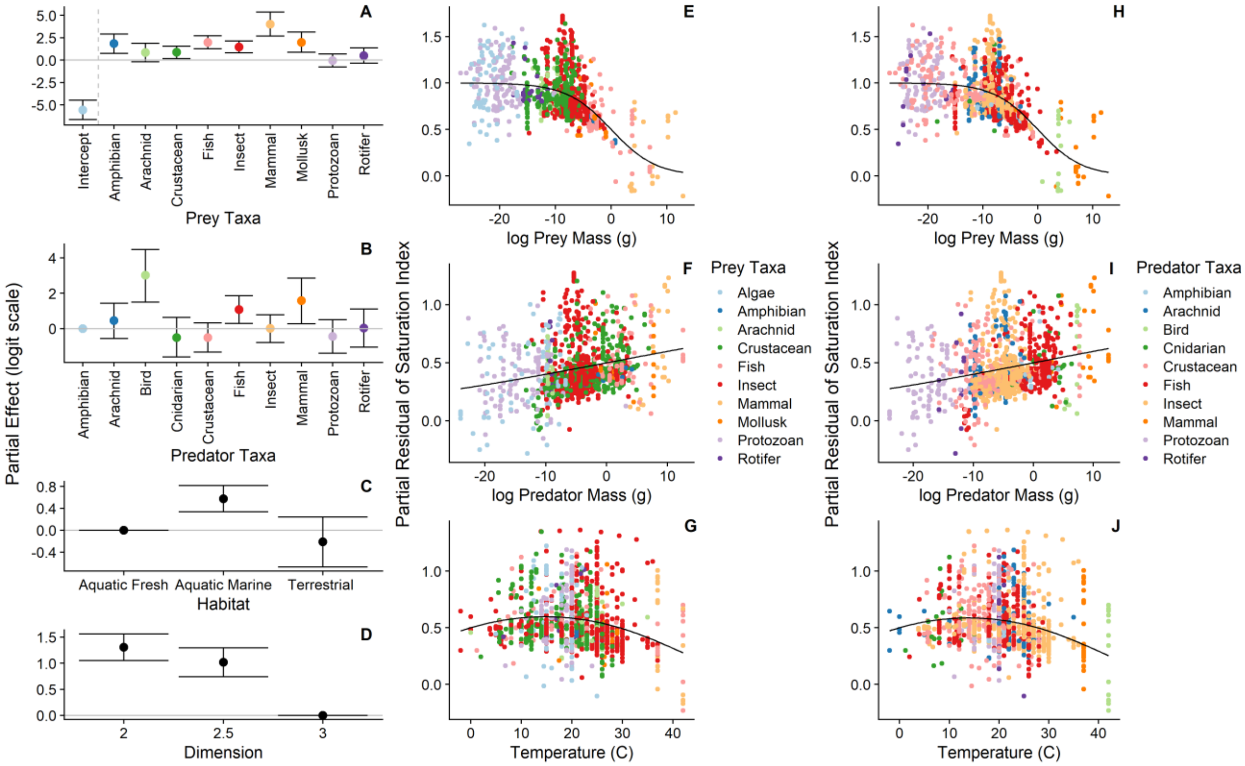


**Figure S5.1. For the model in including birds and mammals, prey taxa (A), predator taxa (B), habitat (C), and dimension (D) exhibited partial effects on the saturation index on the logit scale (the error bars represent 90% credible intervals). The saturation index decreased with prey mass (E,H), increased with predator mass (F,I), and showed a unimodal, concave relationship with temperature (G,J). Note that E-G and H-J include the same data, but E-G are color-coded by prey taxa and H-J are color-coded by predator taxa.**

**Table S5.1.** The estimated parameters and 90% credible intervals (CrI) for each term in the model of the effects of covariates on the index of saturation at high predicted prey densities.

| Model Term | Estimate | 90% CrI |
|---|---|---|
| **Fixed Effects** | | |
| Intercept (Corresponds to an Amphibian feeding on Algae in freshwater in three dimensions) | -5.57 | (-6.66, -4.49) |
| *Prey Major Taxa Effects* | | |
| Amphibian | 1.86 | (0.76, 2.92) |
| Arachnid | 0.84 | (-0.18, 1.88) |
| Crustacean | 0.86 | (0.16, 1.57) |
| Fish | 1.98 | (1.27, 2.73) |
| Insect | 1.46 | (0.81, 2.13) |
| Mammal | 4.01 | (2.69, 5.36) |
| Mollusk | 1.98 | (0.87, 3.14) |
| Protist | -0.04 | (-0.76, 0.68) |
| Rotifer | 0.49 | (-0.35, 1.36) |
| *Predator Major Taxa Effects* | | |
| Arachnid | 0.45 | (-0.55, 1.42) |
| Bird | 3.02 | (1.50, 4.47) |
| Cnidarian | -0.51 | (-1.61, 0.63) |
| Crustacean | -0.51 | (-1.33, 0.32) |
| Fish | 1.07 | (0.29, 1.86) |
| Insect | 0.01 | (-0.80, 0.78) |
| Mammal | 1.58 | (0.27, 2.86) |
| Protist | -0.44 | (-1.39, 0.50) |
| Rotifer | 0.03 | (-1.05, 1.10) |
| *Habitat Effects* | | |
| Aquatic-Marine | 0.58 | (0.34, 0.81) |
| Terrestrial | -0.21 | (-0.67, 0.24) |
| *Dimension Effects* | | |
| 2D | 1.31 | (1.05, 1.56) |
| 2.5D | 1.02 | (0.74, 1.29) |
| *Continuous Variables* | | |
| log Prey Mass (g) | -0.25 | (-0.28, -0.22) |
| log Predator Mass (g) | 0.04 | (0.01, 0.07) |
| Temperature (C) | 0.05 | (0.01, 0.09) |
| Temperature$^2$ | -0.0017 | (-0.003, -0.001) |
| **Random Effects** | | |
| Prey Minor Taxa Standard Deviation | 0.55 | (0.38, 0.76) |
| Predator Minor Taxa Standard Deviation | 0.49 | (0.29, 0.76) |
| **Distribution Parameters** | | |
| Beta Distribution Phi | 2.86 | (2.64, 3.08) |

### S6 Analysis of Space Clearance Rate and Handling Time Covariates

Here we provide tables including the estimates of the regression coefficients of the models examining the relationships between space clearance rates and handling times and their covariates.

**Table S6.1.** The estimated parameters and 90% credible intervals (CrI) for each term in the model of the effects of covariates on the space clearance rates.

| Model Term | Estimate | 90% CrI |
|---|---|---|
| **Fixed Effects** | | |
| Intercept (Corresponds to an Amphibian feeding on Algae in freshwater in three dimensions) | -8.5 | (-10.9, -6.2) |
| *Prey Major Taxa Effects* | | |
| Amphibian | 1.66 | (-0.52, 3.97) |
| Arachnid | 2.13 | (0.015, 4.29) |
| Crustacean | 1.49 | (-0.02, 2.92) |
| Fish | 2.15 | (0.71, 3.64) |
| Insect | 2.71 | (1.24, 4.12) |
| Mollusk | 1.1 | (-1.19, 3.36) |
| Protist | -0.03 | (-1.56, 1.45) |
| Rotifer | 2.37 | (0.59, 4.09) |
| *Predator Major Taxa Effects* | | |
| Arachnid | -1.55 | (-4.03, 1.1) |
| Cnidarian | -0.15 | (-2.98, 2.69) |
| Crustacean | -2.96 | (-5.09, -0.84) |
| Fish | 3.87 | (1.86, 5.9) |
| Insect | -1.18 | (-3.21, 0.92) |
| Protist | -6.63 | (-9.0, -4.27) |
| Rotifer | -2.63 | (-5.35, 0.06) |
| *Habitat Effects* | | |
| Aquatic-Marine | 0.78 | (0.29, 1.25) |
| Terrestrial | -2.71 | (-3.63, -1.82) |
| *Dimension Effects* | | |
| 2D | 4.61 | (4.08, 5.15) |
| 2.5D | 3.99 | (3.43, 4.55) |
| *Continuous Variables* | | |
| log Prey Mass (g) | 0.04 | (-0.016, 0.09) |
| log Predator Mass (g) | 0.35 | (0.3, 0.4) |
| Temperature (C) | 0.24 | (0.17, 0.31) |
| Temperature$^2$ | -0.01 | (-0.007, -0.003) |
| Arena Size | 0.02 | (-0.057, 0.09) |
| **Random Effects** | | |
| Prey Minor Taxa Standard Deviation | 1.19 | (0.89, 1.54) |
| Predator Minor Taxa Standard Deviation | 1.45 | (1.11, 1.86) |
| **Distribution Parameters** | | |
| Residual Standard Deviation | 1.95 | (1.89, 2.02) |

**Table S6.2.** The estimated parameters and 90% credible intervals (CrI) for each term in the model of the effects of covariates on handling times.

| **Model Term** | **Estimate** | **90% CrI** |
|---|---|---|
| **Fixed Effects** | | |
| Intercept<br>(Corresponds to an Amphibian feeding on Algae in freshwater in three dimensions) | -10.91 | (-12.97, -8.88) |
| *Prey Major Taxa Effects* | | |
| Amphibian | 9.3 | (7.33, 11.23) |
| Arachnid | 6.4 | (4.44, 8.3) |
| Crustacean | 6.95 | (5.69, 8.26) |
| Fish | 8.93 | (7.67, 10.23) |
| Insect | 6.71 | (5.5, 7.95) |
| Mollusk | 9.79 | (7.85, 11.72) |
| Protist | 2.26 | (0.95, 3.6) |
| Rotifer | 5.98 | (4.48, 7.57) |
| *Predator Major Taxa Effects* | | |
| Arachnid | 1.71 | (-0.52, 3.94) |
| Cnidarian | -1.25 | (-3.62, 1.08) |
| Crustacean | 1.38 | (-0.37, 3.13) |
| Fish | -1.86 | (-3.6, -0.12) |
| Insect | 0.85 | (-0.92, 2.57) |
| Protist | 5.96 | (3.66, 7.74) |
| Rotifer | 2.16 | (-0.01, 4.42) |
| *Habitat Effects* | | |
| Aquatic-Marine | 0.41 | (0.04, 0.77) |
| Terrestrial | 2.5 | (1.78, 3.2) |
| *Dimension Effects* | | |
| 2D | -1.09 | (-1.5, -0.67) |
| 2.5D | -1.3 | (-1.73, -0.89) |
| *Continuous Variables* | | |
| log Prey Mass (g) | 0.14 | (0.10, 0.18) |
| log Predator Mass (g) | -0.24 | (-0.28, -0.2) |
| Temperature (C) | -0.1 | (-0.16, -0.05) |
| Temperature$^2$ | 0.001 | (-0.00033, 0.002) |
| Arena Size | 0.1 | (0.04, 0.15) |
| **Random Effects** | | |
| Prey Minor Taxa Standard Deviation | 1.15 | (0.89, 1.45) |
| Predator Minor Taxa Standard Deviation | 1.25 | (0.95, 1.65) |
| **Distribution Parameters** | | |
| Residual Standard Deviation | 1.45 | (1.4, 1.49) |

### S7 Generalization of Results to Common Functional Response Forms

Our results in the main text focus on the saturation of feeding rates for a Type II predator functional response. However, a variety of other functional responses are common including functional responses that describe predator feeding rates when the predator's space clearance rate is a function of prey density (a Type III functional response), when predator interference leads to reduction in feeding rates with predator densities, when predators feed on multiple prey types, and more. Here we show for a general functional response form that the saturation of feeding rates with respect to a focal prey density has a similar form as our saturation index for the Type II functional response. We also derive saturation indices for the Type III functional response, the Beddington-DeAngelis functional response, and the multispecies Type II functional response. These indices show that our estimates of feeding rate saturation may be an overestimate of saturation for other functional response forms and support our general conclusion that predator feeding rates may generally be unsaturated under typical prey densities.

#### Saturation Index for a Generalized Functional Response

We first derive the saturation index for a generalized functional response form. For this generalized functional response, we assume that 1) the predator shows saturation with respect to the density of a focal prey species, 2) the space clearance rate can be a function of prey density which can lead to a Type III functional response, and 3) the denominator of the functional response on the focal prey can also include the dependence of predator feeding rates on the densities of alternative prey or the density of the predator. This generalized functional response form is

$$f_G(\mathrm{R}, x) = \frac{\alpha h \mathrm{R}^{q}}{1+\alpha h \mathrm{R}^{q}+g(x)} \qquad \text{eqn. S7.1}$$

where R is the prey density, α is a parameter that gives the space clearance rate when the prey abundance is equal to one, $q$ is a value greater than one that represents the dependence of the space clearance rate on prey density as in a Type III functional response (Holling 1959; Real 1977), $h$ is the handling time of the focal prey, and $g(x)$ represents additional terms in the denominator such as the reduction in the feeding rate due to the handling of alternative prey or the reduction of the feeding rate with predator densities due to predator interference. We can develop an index of feeding rate saturation for this functional response form similar to the index for the Type II functional response in the main text by considering the reduction in feeding rate between a version of $f_G(\mathrm{R}, x)$ that does not saturate with the density of the focal prey ($f_{G-S}(\mathrm{R}, x)$) and $f_G(\mathrm{R}, x)$. In this case, we have

$$f_{G-S}(\mathrm{R}, x) = \frac{\alpha \mathrm{R}^{q}}{1+g(x)} \qquad \text{eqn. S7.2}$$

And the saturation index is given by

$$I = \frac{f_{G-S}(\mathrm{R},x) - f_G(\mathrm{R},x)}{f_{G-S}(\mathrm{R},x)} \qquad \text{eqn. S7.3}$$

Plugging $f_G(\mathrm{R}, x)$ and $f_{G-S}(\mathrm{R}, x)$ into the formula for $I$ gives

$$I = \frac{\frac{\alpha \mathrm{R}^{q}}{1+g(x)} - \frac{\alpha \mathrm{R}^{q}}{1+\alpha h \mathrm{R}^{q}+g(x)}}{\frac{\alpha \mathrm{R}^{q}}{1+g(x)}} \qquad \text{eqn. S7.4}$$

Following steps similar to those in Supplemental Information S1, we get

$$I = 1 - \frac{(\alpha R^q)\left(1 + g(x)\right)}{(\alpha R^q)\left(1 + \alpha h R^q + g(x)\right)}$$

$$= 1 - \frac{1 + g(x)}{1 + \alpha h R^q + g(x)}$$

$$= \frac{1 + \alpha h R^q + g(x)}{1 + \alpha h R^q + g(x)} - \frac{1 + g(x)}{1 + \alpha h R^q + g(x)}$$

$$= \frac{\alpha h R^q}{1 + \alpha h R^q + g(x)} \qquad \text{eqn. S7.5}$$

From this form of the saturation index, we can generalize the results from the analysis in the main text. In terms of the exponent $q$, this parameter only makes the saturation index have a sigmoidal increase with prey density to one rather than a monotonic increase to one. Because feeding rates will be similar in magnitude for both the Type II and Type III functional responses regardless of the shape of the functional response, α values will be generally lower than space clearance rates and predators with a Type III-like functional response will still have feeding rates that saturate at the same level as the Type II functional response given an equal handling time. However, predators with a Type-III-like functional response will be less saturated at lower prey densities due to the sigmoidal approach to saturation at high prey density values. In terms of additional mechanisms appearing in the denominator that reduce predator feeding rates of the functional response such as the handling of non-focal prey and predator interference, these processes will decrease the level of feeding rate saturation at a given focal prey abundance.

Below, we give the specific derivations for the saturation index for three common alternatives to the Type II functional response (a Type III functional response, the Beddington-DeAngelis functional response, and the multispecies Type II functional response) to give concrete examples.

**Saturation Index for the Type III Functional Response**

Type III functional responses describe the case when the predators space clearance rate on the prey is a function of the prey density. In the Type III functional response, we can think of the space clearance rate as being an increasing function of prey density and the Type III functional response $f_{III}$ is given by

$$f_{III} = \frac{\alpha R^q}{1 + \alpha h R^q} \qquad \text{eqn. S7.6}$$

where α is the space clearance rate when the prey density is equal to one, q is a value greater than one representing the dependence of the space clearance rate on prey densities, $h$ is the handling time, and R is the resource density. If the predator feeding rates did not saturate with prey density, the feeding rate of the predator is given by

$$f_{III-S} = \alpha R^q \qquad \text{eqn. S7.7}$$

From equation S7.5, the saturation index for the Type III functional response is then

$$S = \frac{\alpha h R^q}{1 + \alpha h R^q} \qquad \text{eqn. S7.8}$$

As mentioned above, the Type III functional response leads to a sigmoidal increase in saturation with increasing prey densities but otherwise should give similar saturation results to those presented in the main text for the Type II functional response except for a decrease in saturation at low prey densities due to the sigmoidal shape.

**Saturation Index for the Beddington-DeAngelis Functional Response**

The Beddington-DeAngelis functional response is a functional response derived to model the impacts of predator densities on feeding rates through mutual interference (Beddington 1975; DeAngelis *et al.* 1975). The Beddington-DeAngelis functional response $f_{BD}$ is

$$f_{BD} = \frac{a\mathrm{R}}{1+ah\mathrm{R}+\gamma\mathrm{C}} \qquad \text{eqn. S7.9}$$

Where $a$ is the space clearance rate, R is the density of the prey, $h$ is the handling time, γ is the interference rate among predators, and C is the density of predators. If the predator feeding rate did not saturate with prey density, the feeding rate of the predator would be given by

$$f_{BD-S} = \frac{a\mathrm{R}}{1+\gamma\mathrm{C}} \qquad \text{eqn. S7.10}$$

From equation S7.5, the saturation index for the Beddington-DeAngelis functional response is

$$I = \frac{ah\mathrm{R}}{1+ah\mathrm{R}+\gamma\mathrm{C}} \qquad \text{eqn. S7.11}$$

Therefore, predator dependence in the functional response should act to reduce the level of feeding rate saturation at a particular prey density relative to the Type II functional response which does not incorporate predator dependence.

**Saturation Index for the Multispecies Type II functional response**

The multispecies Type II functional response is an extension of the Type II functional response for generalist predators feeding on multiple prey (Murdoch & Oaten 1975; DeLong 2021). The multispecies Type II functional response $f_M$ on a focal prey (chosen without loss of generality to be prey 1) is given by

$$f_M = \frac{a_1\mathrm{R}_1}{1+a_1h_1\mathrm{R}_1+\sum_{i=2}^{S}a_ih_i\mathrm{R}_i} \qquad \text{eqn. S7.12}$$

where $a_j$ is the space clearance rate on prey species $j$, $h_j$ is the handling time on species $j$, $\mathrm{R}_j$ is the density of prey $j$, and $S$ is the total number of prey species in the diet of the predator. If the predator feeding rate did not saturate with respect to the density of the focal prey, the feeding rate of the predator would be given by

$$f_{M-S} = \frac{a_1\mathrm{R}_1}{1+\sum_{i=2}^{S}a_ih_i\mathrm{R}_i} \qquad \text{eqn. S7.13}$$

From equation S7.5, the saturation index for the multispecies functional response with respect to the density of the focal prey is

$$I = \frac{a_1h_1\mathrm{R}_1}{1+a_1h_1\mathrm{R}_1+\sum_{i=2}^{S}a_ih_i\mathrm{R}_i} \qquad \text{eqn. S7.14}$$

Therefore, the additional prey in the diet of the predator and their handling times reduce the level of feeding rate saturation at a particular density relative to the case in which the predator exhibits a Type II functional response and only consumes the focal prey species.

The above result on feeding rate saturation focuses solely on the degree of feeding rate saturation with respect to a focal prey species. However, we can also consider the degree of feeding rate saturation for the total predator feeding rate across all prey. The total feeding rate for the multispecies Type II functional response $f_{MT}$ is

$$f_{MT} = \frac{\sum_{i=1}^{S} a_i \mathrm{R}_i}{1+\sum_{i=1}^{S} a_i h_i \mathrm{R}_i} \qquad \text{eqn. S7.14}$$

Where all parameters are defined above. If the predator's feeding rate did not saturate with respect to any of the prey densities, the feeding rate of the predator would be given by

$$f_{MT-S} = \sum_{i=1}^{S} a_i \mathrm{R}_i \qquad \text{eqn. S7.15}$$

Following algebraic steps similar to those in Supplemental Information S1, we can get the saturation index for the total feeding rate of the predator as

$$I = \frac{\sum_{i=1}^{S} a_i h_i \mathrm{R}_i}{1+\sum_{i=1}^{S} a_i h_i \mathrm{R}_i} \qquad \text{eqn. S7.16}$$

Note that the saturation index for the total feeding rate of the predator is simply the sum of the saturation index with respect to each prey species separately. This means that although additional prey species in a predator's diet decreases the amount of saturation with respect to each prey species separately, increasing the number of prey species in the predator's diet can increase the saturation of the total predator feeding rate. Specifically, since $\lim_{x \to \infty} \frac{x}{1+x} = 1$, if none of the parameters change as additional prey are added to the predator's diet, a multi-prey functional responses will lead to greater saturation of predator feeding rates. However, predator space clearance or handling times may also be lower in the multi-prey case than in the single prey case (Okuyama 2010; Stouffer & Novak 2021), which may make the degree of saturation change with the addition of prey dependent on the degree to which space clearance rates and handling times decrease with additional prey. Unfortunately, little is known about multispecies functional responses and consequently any generalizations are difficult.
